# Naïve orangutans (*Pongo abelii & Pongo pygmaeus*) individually acquire nut-cracking using hammer tools

**DOI:** 10.1101/2020.04.21.052712

**Authors:** Elisa Bandini, Johannes Grossmann, Martina Funk, Anna Albiach Serrano, Claudio Tennie

## Abstract

Nut-cracking using hammer tools has been argued to be one of the most complex tool-use behaviours observed in non-human animals (henceforth: animals). Recently, even the United Nations Convention on the Conservation of Migratory Species (CMS) recognised the unique nature of chimpanzee nut-cracking by making it the first animal behaviour to be awarded UN-protected status (Picheta, 2020). So far, only chimpanzees, capuchins and macaques have been observed using tools to crack nuts in the wild (Boesch & Boesch, 1990; Gumert, Kluck, & Malaivijitnond, 2009; Ottoni & Mannu, 2001). However, the learning mechanisms behind this behaviour, and the extent of nut-cracking in other primate species are still unknown. The aim of this study was two-fold. First, we aimed to examine whether other great ape species would develop nut-cracking when provided with all the tools and motivation to do so. Second, we wanted to examine the mechanisms behind the emergence of nut-cracking in a naïve sample. Orangutans (*Pongo abelii; pygmaeus*) have not been observed cracking nuts in the wild, despite having the second most extensive tool-use repertoire of the great apes (after chimpanzees), having the materials for the behaviour in the wild (albeit rarely) and possessing flexible problem-solving capacities. Therefore, orangutans are a valid candidate species for the investigation of the development of nut-cracking. Four nut-cracking-naïve orangutans at Leipzig zoo (*Pongo abelii*; *M*_*age*_=16; age range=10-19; 4F; at time of testing) were provided with nuts and hammers but were not demonstrated the nut-cracking behavioural form, in order to control for the role of copying social learning in the acquisition of this behaviour. Additionally, we report data from a previously unpublished study by one of the authors (MF) with eight orangutans housed at Zürich zoo (10 *Pongo abelii* and two *Pongo pygmaeus;* M_age_=14; age range =2-30; 5F; at time of testing) that followed a similar testing paradigm. Out of the twelve orangutans across both testing institutions, at least four individuals, one from Leipzig (*Pongo abelii*) and three from Zürich (*Pongo abelii*; *pygmaeus*), spontaneously expressed nut-cracking with a wooden hammer. These results suggest that the behavioural form of nut-cracking using hammer tools can emerge in orangutans when required through individual learning combined, in some cases, with non-copying social learning mechanisms.

## Introduction

Once heralded as the main distinguishing feature of humans in the animal kingdom, it is now known that several other species also possess the ability to use tools (Shumaker, Walkup, Beck, & Burghardt, 2011). Of these species, non-human great apes (henceforth apes), alongside New Caledonian crows (Kenward et al., 2011), demonstrate the most extensive and potentially most ‘complex’ tool-use repertoires (van Schaik, Deaner, & Merrill, 1999), with complex tool-use defined here as: *“tool-use variants that include at least two tool elements (for example, hammer and anvil), flexibility in manufacture or use (that is, tool properties are adjusted to the task at hand), and that skills are acquired in part by social learning”* (Meulman, Sanz, Visalberghi, & van Schaik, 2012, 58). Of the tool-use behaviours observed in wild and captive great apes, nut-cracking using hammer tools (henceforth: nut-cracking) in chimpanzees is often cited as one of the most complex behaviours within their tool-use repertoires (Boesch, 1991; Lonsdorf, 2013). The claims for the complexity of nut-cracking in chimpanzees are based on several assumptions on the behaviour, including the learning mechanisms underlying the acquisition of nut-cracking in wild chimpanzees. Whilst we will discuss these assumptions in further detail in the following section, below we provide definitions for the main social learning mechanisms discussed here.

The animal social learning field is rife with terminology and mechanisms (for an overview of these terms, see Whiten, Horner, Litchfield, & Marshall-Pescini, 2004). However, throughout this paper we will mainly be referring to two types of social learning: copying and non-copying social learning mechanisms. Copying social learning refers to social learning mechanisms that can transmit the actual form of a behaviour and/or artefact (where form is defined as “*the specific action [and/or artefact] component(s) and organisation of a behaviour”;* Bandini & Tennie, 2020, 2). Copying social learning includes mechanisms such as imitation and some types of emulation (Bandini & Tennie, 2020). On the other hand, non-copying social learning mechanisms are those that cannot transmit the form of a behaviour and/or artefact. Instead, these mechanisms can regulate the frequency of the acquisition of a behaviour, for example by increasing an individual’s motivation to interact or manipulate a certain object or location. Non-copying social learning mechanisms include local and/or stimulus enhancement (Bandini & Tennie, 2020).

### Nut-cracking in primates

Nuts represent an important source of calories and fat in primates’ diets (Biro et al., 2003). The encased condition of these nutrients, however, makes it often beneficial and sometimes neccessary to use stone and/or wooden hammers to crack open the nuts against hard surfaces. Nut-cracking has been observed in chimpanzees, long-tailed macaques, and capuchins (Boesch & Boesch, 1990; Gumert, Kluck, & Malaivijitnond, 2009; Ottoni & Mannu, 2001). Of all nut-cracking behaviours, however, the best-studied example is that of chimpanzees (Biro et al., 2003; Boesch & Boesch, 1990; Luncz & Boesch, 2014; Luncz, Mundry, & Boesch, 2012). Indeed, this behaviour in chimpanzees has recently been selected for conservation by the United Nations Convention on the Conservation of Migratory Species (CMS) body, showing how important – and how cultural - this behaviour is considered to be, even by organisations outside of academia (Picheta, 2020).

Wild chimpanzees in the Taï Forest (Ivory Coast) and in Bossou (Guinea) use hammer tools to access the kernels of several nut species - *Panda oleosa, Parinari excelsa, Saccoglottis gabonensis, Coula edulis*, and *Detarium senegalensis* (Proffitt, Haslam, Mercader, Boesch, & Luncz, 2018). . The crux of the nut-cracking behavioural form in chimpanzees (see also Foucart et al., 2005) involves three steps: (1) Retrieving a nut from the surrounding area and placing it on an anvil (e.g., a tree root or a stone), (2) Picking up a stone- or wooden *hammer* (with one hand or both hands) and (3) Hitting the nut with the *hammer* (holding it with one or both hands) until it is open and the inside kernel can be retrieved and consumed (Boesch & Boesch, 1983; Carvalho, Biro, McGrew, & Matsuzawa, 2009). Sometimes more steps are described, such as the transportation of the materials to the nut-cracking site (Carvalho et al., 2009) and the stabilisation of the anvil on the ground (although this is a rare behaviour; Carvalho et al., 2009). This multi-step approach has been regarded as a complex tool use behaviour (Meulman et al., 2012), because it is improbable that such a compound behaviour is acquired in its entirety by chance, especially considering that it is only rewarded at the end of the chain of actions (note that most of the other behavioural forms within chimpanzee tool-use repertoire only involve the manipulation of a single object (usually a stick) and include only one action (e.g., marrow picking; see Whiten et al., 2001 for an overview of chimpanzee behaviours and their description). Moreover, the precision needed to crack open nuts is said to contribute to the complexity of the behaviour since (at least at the beginning) many attempts will go unrewarded. Perhaps for these reasons, chimpanzee nut-cracking is often assumed to be driven and maintained by action copying (Boesch, 1991; Boesch, Marchesi, Marchesi, Fruth, & Joulian, 1994), although this assumption is heavily debated (e.g., see Neadle et al., 2020).

### Acquisition of nut-cracking in chimpanzees

Juvenile chimpanzees take a long time to acquire nut-cracking (Biro et al., 2003; Boesch & Boesch, 1990) and observations of wild juvenile chimpanzees suggest that the acquisition of nut-cracking may occur within a sensitive learning period, most likely when chimpanzees are between the ages of three and five years old (Inoue-Nakamura & Matsuzawa, 1997). If the behaviour is not acquired within this sensitive learning period, chimpanzees will seemingly never develop the behaviour (Biro et al., 2003). This seems to also be the case for nut-cracking in other primates, such as long-tailed macaques (Tan, 2017).

Furthermore, it has been proposed that during this learning period, juveniles acquire nut-cracking by observing and then copying their mother’s actions (e.g., see Biro et al., 2003) and that a repeated cycle of observation and practice sessions is required before nut-cracking can be expressed (e.g., what Whiten, 2017, 7795, describes as a “helical process of learning”). In a similar interpretation, de Waal (2008) suggests that juvenile chimpanzees copy their mothers’ nut-cracking via ‘Bonding and Identification-based Observational Learning’ (BIOL), where a juvenile copies actions in order “to be like others” (de Waal, 2008, 231). Yet, a lengthy learning period alone is not necessarily indicative of copying. Instead, it can be also be explained by mere maturation processes, alongside an extended period of individual learning (likely encouraged by non-copying variants of social learning, such as stimulus and local enhancement; Whiten et al., 2004 and “peering”; Corp & Byrne, 2002; Schuppli et al., 2016). Furthermore, conclusive evidence for (unenculturated) apes possessing the ability to copy behaviours is still lacking. Whilst enculturated and heavily trained primates have demonstrated an ability to copy actions (indeed, this type of training seems to change the individuals’ brain structures to allow action copying; Hecht et al., 2013; Pope, Taglialatela, Skiba, & Hopkins, 2017), experimental paradigms aimed at identifying this ability in unenculturated chimpanzees have so far been unsuccessful (Clay & Tennie, 2017; Tennie, Call, & Tomasello, 2012; Tomasello et al., 1997). Therefore, whilst further studies should be carried out on this ability in different ape species, the current state of knowledge suggests that alternative explanations to those based solely on copying social learning should be explored. One alternative approach towards explaining the acquisition of nut-cracking in primates is provided by the zone of latent solutions hypothesis (ZLS; Tennie, Call, & Tomasello, 2009). The ZLS hypothesis argues that behavioural forms are acquired in many species via an interplay of individual learning and non-copying variants of social learning. According to this hypothesis, the behavioural form of primate nut-cracking would not be copied, but individually derived. There are many ways in which this individual learning may work. To give just one example, the difficulty of learning individually such a complex behaviour may be overcome by individuals having a general predisposition to explore and manipulate objects plus some cognitive capacities like good spatial memory (that allows to locate the needed materials), inhibitory control (that allows to delay a reward), planning abilities and working memory (that allow to chain steps towards a goal) and some understanding of the physical affordances of objects and of object relations (that can aid in the selection of appropriate materials and actions to process the materials). As a result, such individuals are able to solve problems in a flexible way. This would explain why, for example, wild nut-cracking chimpanzees use different types of anvils (stationary and non-stationary) depending on their needs and material availability. The ZLS hypothesis also suggests that the observed differences in nut-cracking activity across chimpanzee populations are fostered by non-copying social learning mechanisms (such as local and/or stimulus enhancement), which increase the likelihood of reinnovation once a population already contains individuals who have innovated the behavioural form. This can then lead to a frequency increase and maintenance of the behavioural forms in question in some populations but not in others. The ZLS hypothesis predicts successful reinnovation of behavioural forms by naïve ape subjects provided the right conditions and in the absence of copying opportunities. This prediction has been met several times, as demonstrated by a fast-growing experimental literature detailing successful individual acquisitions of various wild-type behavioural forms (including tool use) in various species of naïve, captive great apes (Allritz, Tennie, & Call, 2013; Bandini & Tennie, 2017; 2019; 2020; Bandini & Harrison, 2020; Menzel, Fowler, Tennie, & Call, 2013; Neadle, Allritz, & Tennie, 2017; Tennie, Hedwig, Call, & Tomasello, 2008). Therefore, the ape ZLS hypothesis has growing support, but whether it can also explain the behavioural form of nut-cracking is still an open question.

### Latent solutions testing methodology

In order to examine whether a subject can acquire a behavioural form and the mechanisms behind this acquisition (whether it is driven by copying social learning or by individual learning and non-copying social learning), Tennie & Hedwig (2009) described the ‘latent solutions’ (LS) testing methodology. This methodology allows for the role of individual learning in the acquisition of a target behavioural form. All the ecological materials of the target behavioural form, but no demonstrations, are provided to naïve subjects, who have never seen, or been trained in, the target behaviour before. Subjects should be so-called ‘ecologically-representative’ individuals, i.e. unenculturated captive animals who live in social groups (Henrich & Tennie, 2017). If the target behavioural form emerges under these conditions, then, logically, it can be concluded that copying is not *required* for the form of behaviour to emerge. If the behaviour does not emerge in this baseline condition, then it could be that some variant of copying is necessary for the behaviour to be acquired (for these cases, Bandini & Tennie, 2018 provide an extended LS testing methodology that allows for the level and variant of social learning required (if any) to be identified), *or* that other factors, such as sensitive periods, opportunities to practice or motivation levels, play a role (Bandini & Tennie, 2018; Neadle et al., 2020). Past LS studies have demonstrated that multiple behavioural forms – including tool use behavioural forms – can be individually acquired by primates (see above). For example, naïve, captive, capuchins were tested following the LS testing methodology and two individuals spontaneously started cracking nuts, without copying the behaviour (Visalberghi, 1987). Furthermore, this methodology has also shown that different species may sometimes overlap in their latent solution repertoires. Indeed, human children were recently found to spontaneously acquire various tool-use behaviours observed in wild chimpanzees and orangutans (Reindl, Beck, Apperly, & Tennie, 2016, Nelder et al., 2020; and see also: Allritz et al., 2013; Bandini & Tennie, 2019, 2017; Menzel et al., 2013; Neadle et al., 2017; Tennie et al., 2008 for examples of latent solutions across primate species).

With regard to great apes, apart from the observations of chimpanzee nut-cracking, anecdotal reports exist of gorillas and bonobos cracking nuts in sanctuaries (Wrangham, 2006) – though the exact circumstances of these innovations remain unclear, whilst orangutans have not (perhaps not yet) been reported to crack nuts (Fox et al., 2004; Parker & Gibson, 1977). Yet, despite being the most arboreal great apes, orangutans have the second most extensive tool-use repertoire in the wild after chimpanzees (Fox, van Schaik, Sitompul, & Wright, 2004; Meulman & van Schaik, 2013). Orangutans’ tool-use repertoires include extractive foraging tool-use and, similarly to that of chimpanzees, demonstrate intra and inter-population variation (van Schaik & Knott, 2001). Moreover, there are some reports of orangutans using sticks to ‘hammer’ open termite or bee nests (Fox, Sitompul, & van Schaik, 1999). These behaviours are not fully comparable to chimpanzee nut-cracking, as they generally involve thin sticks (whilst chimpanzee wooden hammers are usually thicker and/or heavier than the sticks they use for other tool-use behaviours) and the hammering action used by orangutans is unlikely to carry enough force to break an object as sturdy as an encased nut (Fox et al., 1999). Therefore, orangutans present a promising candidate to acquire nut-cracking, whilst still remaining naïve to this form of hammering behaviour..

Four naïve captive orangutans housed at Leipzig zoo (*M*_*age*_=16; age range=10-19; 4F; at time of testing) were provided with all the raw materials necessary for nut-cracking (nuts, wooden hammers, cracking locations), but with no information or demonstrations on *how* to crack nuts. This was to test whether orangutans could individually acquire this behavioural form of nut-cracking, without having access to a model to copy. After conducting this study, it was brought to the authors’ attention that an unpublished thesis contained a similar study with orangutans, carried out by Dr. Funk (one of the co-authors of the current manuscript) between 1983-1984. In this study, eight naïve captive orangutans housed at Zürich zoo (10 *Pongo abelii* and two *Pongo pygmaeus;* M_age_=14; age range =2-30; 5F; at time of testing) were tested, following a similar (baseline) paradigm as our study. Therefore, the methods and findings from this previous, unpublished, study are also described here. For the sake of clarity, we will refer to Leipzig study and Zürich study in each case.

## Leipzig study

### Methods

#### Ethical approval

In accordance with ethical recommendations, all subjects were housed in indoor and outdoor enclosures containing climbing structures and natural features. Subjects received their regularly scheduled feedings and had access to enrichment devices and water *ad lib*. Subjects were never food or water deprived for the purposes of this study. All research was conducted in the subjects sleeping rooms. An internal committee of the Max Planck Institute for Evolutionary Anthropology (director, research coordinator) and the Leipzig zoo (head keeper, curator, vet) granted ethical approval for this project. No medical, toxicological or neurobiological research of any kind is conducted at the WKPRC. This research was non-invasive and strictly adhered to the legal requirements of Germany. Animal husbandry and research comply with the “EAZA Minimum Standards for the Accommodation and Care of Animals in Zoos and Aquaria”, the “WAZA Ethical Guidelines for the Conduct of Research on Animals by Zoos and Aquariums” and the “Guidelines for the Treatment of Animals in Behavioral Research and Teaching” of the Association for the Study of Animal Behavior (ASAB).

#### Subjects

Research was carried out at the Wolfgang Köhler Primate Research Center (WKPRC), Leipzig, Germany in 2007. Four orangutans (*M*_*age*_=16; age range=10-19; 4F; at time of testing) participated in the study (see the demographic information in Table 1; all subjects were born (except for DK) and raised at the testing institution). The keepers confirmed that none of the individuals in this study had prior experience with macadamia nuts. Hazelnuts and walnuts, however, had occasionally been provided by the keepers. Yet, the orangutans either opened these with their teeth or, occasionally, by hitting them with their hand against a hard surface. Crucially, none of the orangutans at the WKPRC had ever been observed using a *tool* for nut-cracking before this study. Indeed, heavy objects that could potentially be used as hammers (such as stones or wooden stumps) are not allowed inside the enclosures of the WKPRC, for health and safety reasons, and therefore the subjects can confidently be assumed to have been naïve to the target behaviour prior to this study.

**Table 1:** Demographic information on the subjects included in the Leipzig zoo study.

| Name | Species | Sex | Date of birth | Place of birth | Breeding |
| --- | --- | --- | --- | --- | --- |
| Dokana (DK) | *Pongo abelii* | F | 31/01/1989 | Dresden, DE | Parent |
| Padana (PD) | *Pongo abelii* | F | 18/11/1997 | Leipzig, DE | Parent |
| Pini (PI) | *Pongo abelii* | F | 30/06/1988 | Leipzig, DE | Parent |
| Dunja (DJ) | *Pongo abelii* | F | 19/04/1990 | Leipzig, DE | Hand |

#### Procedure

We implemented three conditions sequentially (see Table 2): The first condition was a baseline, in which subjects could only acquire the nut-cracking behaviour individually, as no information on the actions required for the behaviour was provided. The second condition was another baseline, which we called locked-anvil condition, that guaranteed that the object provided as an anvil could *only* be used as an anvil and not as a hammer (see below). The third condition was a demonstration condition, in which subjects could potentially learn nut-cracking behaviour through social learning (of any variant) after observing a conspecific model (PD; age 10 at the time of testing). Subjects were tested separately with no visual or acoustic access to each other. While the sub-adult (PD) was tested alone, the adult females were tested together with their dependent offspring (however no data were analysed from the behaviour of the offspring as they were too young at the time of testing to attempt performing the task).

**Table 2:** Table showing the number of conditions, sessions and role of each subject from the Leipzig zoo study.

| Subject | Conditions participated in | Number of sessions per condition | Role |
| --- | --- | --- | --- |
| DK | Baseline, Demonstration | Baseline: 5<br>Demonstration: 5 | Subject<br>Subject |
| PD | Baseline, Locked-anvil, Demonstration | Baseline: 5<br>Locked-anvil: 4<br>Demonstration: 5 | Subject<br>Subject<br>Conspecific model in the demonstration condition |
| PI | Baseline, Demonstration | Baseline: 5<br>Demonstration: 5 | Subject<br>Subject |
| DJ | Baseline, Demonstration | Baseline: 5<br>Demonstration: 5 | Subject<br>Subject |

#### Baseline condition

During each of five baseline sessions, subjects had access to one large wooden block (the anvil; height 30 cm, diameter 50 cm, approximate weight 50 kg) with 5 depressions (diameter 2.5 cm) carved into the top side to facilitate the placement of the nuts, mirroring similar depressions of anvils in the wild (e.g., Carvalho et al., 2009; Luncz et al., 2012), two smaller wooden blocks (the wooden hammers; height 30 cm, diameter 50 cm, approximate weight 2.4 kg each) and five macadamia nuts (see figure 1 below). The materials were scattered evenly on the floor in the testing room, which was emptied of any other objects prior to the test to avoid distractions, within approx. one square meter. The subjects were not allowed to enter the room until all the materials were in place. Sessions lasted a maximum of twenty minutes but were discontinued earlier if the subjects had successfully opened all the nuts. The shells of the opened nuts and any nuts that the subjects did not open were retrieved after each session and discarded. A video camera were used to record the subjects’ behaviour. For each subject, the between-session interval was at least 24 hours.

**Figure 1:** Photograph of the testing apparatus with the anvil, wooden hammers and macadamia nuts in the Leipzig zoo study.

#### Locked-anvil condition

After the baseline condition, the single successful subject (PD, see the results section) participated in four additional sessions that were similar to the initial baseline sessions but with the anvil fixed on the ground (by being pressed down with a sliding door). This way, we encouraged the subject to explore options other than using the anvil as a hammer to crack open the nuts (as in the baseline the subject used an anvil-dropping and rolling technique to crack the nuts).

#### Demonstration condition

After the baseline and locked-anvil conditions, the three orangutans that did not demonstrate the nut-cracking behaviour in the baseline condition participated in five subsequent demonstration condition sessions (15 sessions in total). Before each session, PD, who had reliably started the nut-cracking behaviour in the previous phases, served as a demonstrator, cracking (and eating) five macadamia nuts. The subject, who had access to two hammers and a fixed anvil, could observe PD’s performance from an adjacent cage. As soon as the subject had observed at least one successful nut-cracking bout (coded when the subject had its head oriented towards the demonstrator and its eyes were open during a successful nut-cracking bout by the demonstrator), five macadamia nuts were placed into the subject’s enclosure and the session started. The demonstrations continued even after the nuts were placed in the enclosure. The rest of the testing procedure remained the same as in the baseline condition (see above).

#### Data collection and reliability

We live and video coded the behaviour performed by subjects to try to open the macadamia nuts (see Tables 3 & 4). Two second coders, who were not familiar with the aims and results of the study, watched the videos and coded the same categories as the original coder to assess inter-rater reliability. One coded the ethogram of behaviours, and how often each individual demonstrated the methods, whilst the other coded the number of successes in opening the nuts and the time spent with a nut in the subject’s mouth. A Cohens kappa was calculated to assess the inter-rater reliability of both sets of data. All data are available in OSF (please see: https://osf.io/43fbr/?view_only=fd9290ce18b542c7a43a102f600ab22d).

**Table 3:**
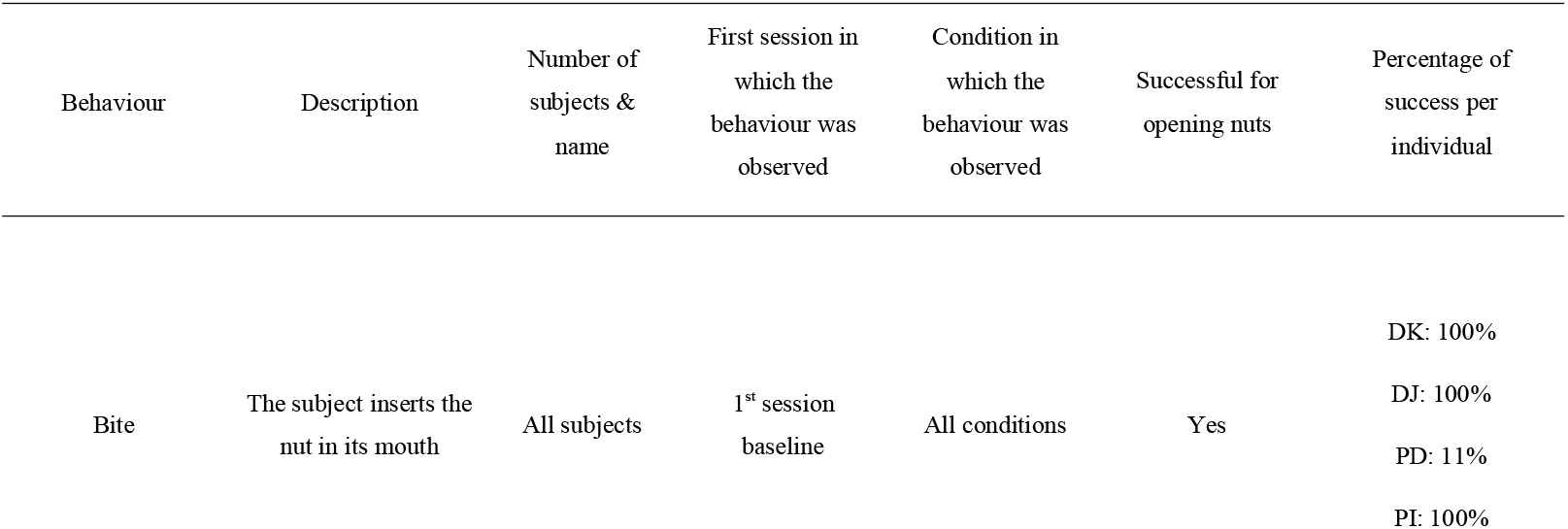
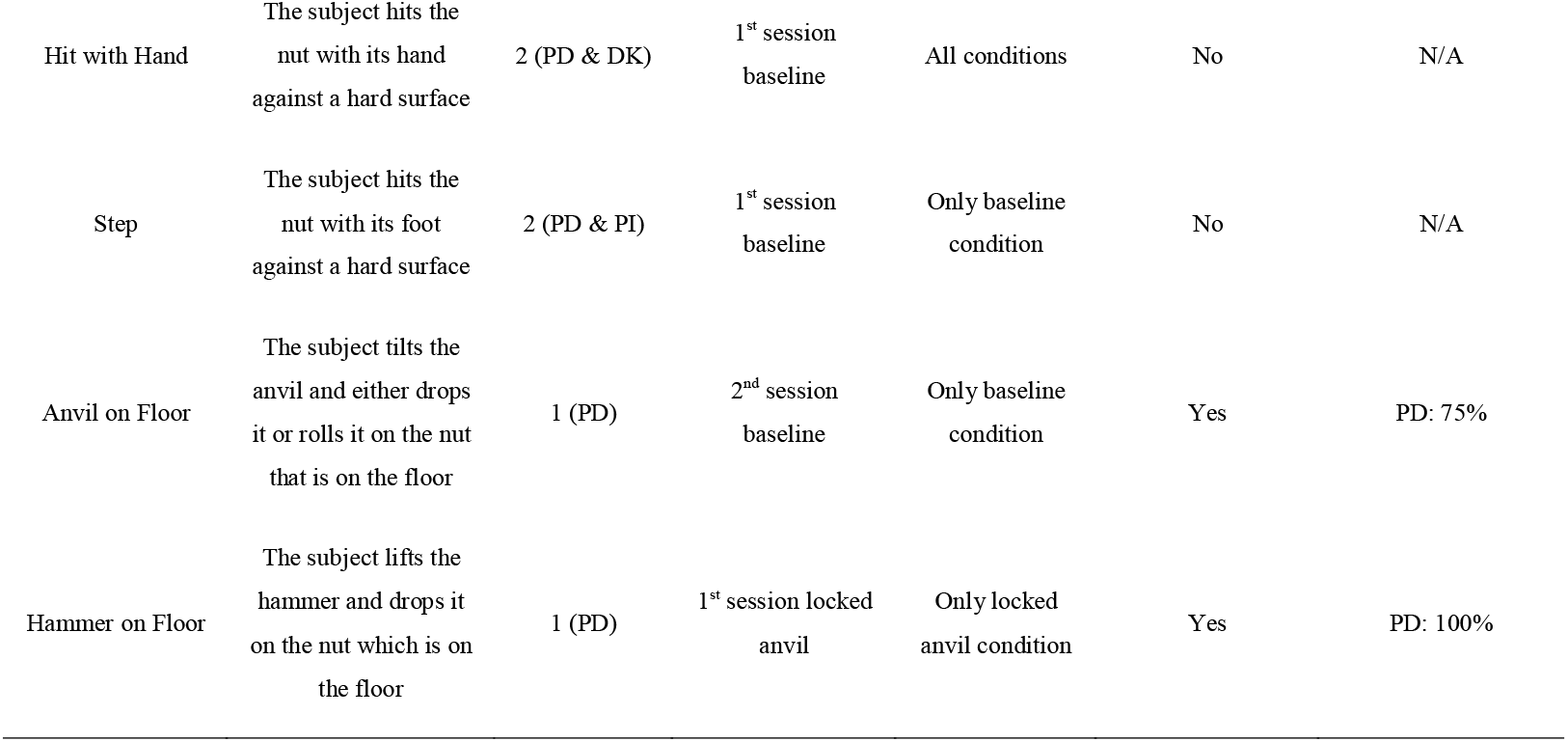
Ethogram of behaviours directed towards the nuts by subjects in the Leipzig zoo study across all conditions.

| Behaviour | Description | Number of subjects & name | First session in which the behaviour was observed | Condition in which the behaviour was observed | Successful for opening nuts | Percentage of success per individual |
| --- | --- | --- | --- | --- | --- | --- |
| Bite | The subject inserts the nut in its mouth | All subjects | 1<sup>st</sup> session baseline | All conditions | Yes | DK: 100%<br>DJ: 100%<br>PD: 11%<br>PI: 100% |
| Hit with Hand | The subject hits the nut with its hand against a hard surface | 2 (PD & DK) | 1<sup>st</sup> session baseline | All conditions | No | N/A |
| Step | The subject hits the nut with its foot against a hard surface | 2 (PD & PI) | 1<sup>st</sup> session baseline | Only baseline condition | No | N/A |
| Anvil on Floor | The subject tilts the anvil and either drops it or rolls it on the nut that is on the floor | 1 (PD) | 2<sup>nd</sup> session baseline | Only baseline condition | Yes | PD: 75% |
| Hammer on Floor | The subject lifts the hammer and drops it on the nut which is on the floor | 1 (PD) | 1<sup>st</sup> session locked anvil | Only locked anvil condition | Yes | PD: 100% |

**Table 4:** Count and percentage of each behaviour directed towards the nuts per individual per condition and sessions in the Leipzig zoo study

| Condition | Subject | Number of sessions | Number of sessions in which Bite was observed | % | Number of sessions in which Hit with Hand was observed | % | Number of sessions in which Step was observed | % | Number of sessions in which Anvil on Floor was observed | % | Number of sessions in which Hammer on Floor was observed | % |
| --- | --- | --- | --- | --- | --- | --- | --- | --- | --- | --- | --- | --- |
| Baseline | DK | 5 | 5 | 100% | 3 | 60% | 0 | 0% | 0 | 0% | 0 | 0% |
|  | DJ | 5 | 5 | 100% | 0 | 0% | 0 | 0% | 0 | 0% | 0 | 0% |
|  | PD | 5 | 5 | 100% | 3 | 60% | 2 | 40% | 4 | 80% | 0 | 0% |
|  | PI | 5 | 5 | 100% | 0 | 0% | 1 | 20% | 0 | 0% | 0 | 0% |
| Baseline Total |  | 20 | 20 | 100% | 6 | 30% | 3 | 15% | 4 | 20% | 0 | 0% |
| Anvil Locked | PD | 4 | 4 | 100% | 2 | 50% | 0 | 0% | 0 | 0% | 4 | 100% |
| Anvil Locked Total |  | 4 | 4 | 100% | 2 | 50% | 0 | 0% | 0 | 0% | 4 | 100% |
| Demonstration | DK | 5 | 5 | 100% | 0 | 0% | 0 | 0% | 0 | 0% | 0 | 0% |
|  | DJ | 5 | 5 | 100% | 2 | 40% | 0 | 0% | 0 | 0% | 0 | 0% |
|  | PI | 5 | 5 | 100% | 0 | 0% | 0 | 0% | 0 | 0% | 0 | 0% |
| Demo Total |  | 15 | 15 | 100% | 2 | 13% | 0 | 0% | 0 | 0% | 0 | 0% |

#### Reliability testing results

A Cohen’s kappa was run to assess the reliability of the coded data. We did not expect to find a very high inter-observer reliability for some of the behaviours, as the data were collected in the orangutans’ management areas (due to the testing facilities requirements), which were dark and often did not allow for a clear view from the filming platform (also as subjects moved dynamically and often blocked camera angles with their bodies). Despite this, in terms of the general coding of the ethogram a moderate agreement (Cohen, 1968) was found (*k*=0.60). However, note that a substantial interobserver agreement (*k*=0.80) was found for our main experimental outcome, namely for coding the target nut-cracking behavioural forms (anvil on floor and hammer on floor behaviours shown by PD). Therefore, whilst we acknowledge that the reliability scores are lower than ideal for some additional behaviours coded in the ethogram, the main behavioural outcome (nut-cracking) was reliably coded. The overarching aim of this study was to examine whether nut-cracking with a tool would develop in naïve orangutans, and these observations received a substantial agreement across coders (*k*=0.80).

## Results

Table 3 presents the behaviours coded, their descriptions, how many individuals attempted each behaviour, the first session in which behaviours were observed, in which experimental conditions they were observed, whether or not they allowed opening nuts and the percentage of times each behaviour resulted in successfully cracking open a nut (see supplementary material for video clips of the most common behaviours observed). In the baseline condition, the juvenile individual, PD (F, 10 years old at time of testing, mother-reared and born at the testing institution; see Table 1), successfully cracked nuts by using the large wooden anvil-block as a hammer-tool (see also Table 3). When, in the locked-anvil condition, the large anvil-block was fixed to the ground and therefore could no longer be used as a hammer, this same subject cracked nuts by using the wooden hammers (see supplementary videos), showing a similar behaviour to chimpanzee nut-cracking. No other individual in the study performed the nut-cracking behaviour with a tool. Instead, the other (all adult) subjects opened the nuts with their teeth (*bite*, see Tables 3 & 4). This *bite* behaviour of adults continued even after the demonstration condition, in which the adults had the opportunity to observe PD cracking nuts using the target behaviour. Indeed, the adults used primarily the *bite* method, followed by the only other method they used: *hit with hand* (see more below).

### Baseline condition

The *bite* method was the first method attempted in the baseline by all individuals, and the one used in a higher number of sessions in this phase (*bite* was attempted in 20/20 (100%) of the sessions, followed by *hit with hand* (6/20, 30%;), *anvil on floor* (4/20, 20%) and *step* (3/20, 15%;). All subjects attempted to open at least some nuts with their mouth, feet or hands in most sessions, whereas only PD used the *anvil on floor* method, in 4/5, 80% of her sessions (from the 2^nd^ session of the baseline). Of these methods, only the *bite* and *anvil on floor* led to successful kernel access. The adult females accessed an average of 4.4 out of 5 nut kernels per session using the *bite* method (and were successful from the first session). PD also tried to open nuts first with the mouth in her first session but failed to open them. However, in the second and third sessions, PD tilted the large wooden block, placed a nut under the block, and then dropped it on the nut. By using this method (*anvil on floor*), PD successfully opened six nuts overall (the remaining nuts stayed unopened, as PD then reverted to attempting the *bite* methodology unsuccessfully). In the fourth session, PD successfully cracked one nut with her mouth but failed to open more nuts with either the *bite* or *anvil on floor* techniques. These data may suggest that PD was relatively incapable of cracking open the nuts with her teeth. In the last session, PD opened all five nuts using the *anvil on floor* method.

### Anvil-locked condition (note: only PD was tested)

This condition (4 sessions) was carried out to examine whether PD would be able to change from her technique of using the large wooden block (which had been devised as an anvil) to using the smaller wooden pieces provided (which were designed to resemble the hammers used by wild chimpanzees). From the first session, PD used the wooden hammers to perform the target nut-cracking behaviour, albeit ignoring the large block as an anvil. Instead, PD placed nuts on the floor (which was sufficiently hard), and then used the wooden hammer to forcibly hit the nut until it opened (i.e., *hammer on floor* occurred in all sessions). Other than *hammer on floor*, only *bite* and *hit with hand* were recorded in this condition. PD cracked 19 of 20 nuts using the *hammer on floor* method and no nuts using the *bite* method.

### Demonstration condition

Despite being provided with live demonstrations from PD of the target nut-cracking behaviour in the demonstration condition (15 sessions in total), none of the adult females subsequently used any of the provided tools to open nuts. All adults continued to crack the nuts using their teeth or by trying to open the nuts (unsuccessfully) using the *hit with hand* method (*bite* 100%, 15/15 of the sessions; *hit with hand* 13%, 2/15 of the sessions). All the nuts that were opened in the demonstration condition were opened with the *bite* behaviour.

## Zürich study

### Methods

#### Ethical approval

No specific ethical protocols were required when this study was carried out (in the 1980s). However, the experiments were non-invasive, and were designed alongside the animals’ keepers to ensure the well-being of the animals. The study strictly adhered to the legal requirements of Switzerland. All subjects were housed in indoor and outdoor enclosures containing climbing structures and natural features. Subjects received their regularly scheduled feedings and had access to enrichment devices and water *ad lib*. Subjects were never food or water deprived for the purposes of this study. The research was conducted in the subjects’ familiar environment and any kind of stress to the animals was avoided.

#### Subjects

Research was carried out at Zürich zoo, Switzerland, between 1983-1984 by MF. Twelve orangutans were tested (10 *Pongo abelii* and two *Pongo pygmaeus;* M_age_=14; age range= 2-30; 5F; see demographic information in Table 6). The keepers confirmed that none of the subjects had any previous experience with coula nuts or Brazil nuts. However, the orangutans had occasionally been provided with walnuts and coconuts. Whilst the walnuts could be easily opened by biting, the coconuts were opened by hitting them against the hard floor or walls of the enclosure. Despite the potential utility of a tool to open the coconuts, the keepers had never observed the orangutans using hammer tools to open the coconuts and had never demonstrated the hammer technique to them – for these or any other nuts. The two *Pongo pygmaeus* were tested together, and the remaining subjects (all the *Pongo abelii*) were tested in two groups (see Table 6 for a list of individuals in each group). These two groups did not have visual access to each other.

**Table 6:** Demographic information on the subjects included in the Zürich zoo study.

| Name | Species | Testing Group | Sex | Date of birth | Place of birth | Breeding |
| --- | --- | --- | --- | --- | --- | --- |
| Rosa (RS) | *Pongo pygmaeus* | 1 | F | 1958 | Wild Caught | Parent |
| Adam (AD) | *Pongo pygmaeus* | 1 | M | 1953 | Wild Caught | Parent |
| Pongo (PG) | *Pongo abelii* | 2 | M | 1961 | Wild Caught | Parent |
| Lea (LA) | *Pongo abelii* | 2 | F | 22/08/1967 | Zürich, CH | Hand |
| Farida (FA) | *Pongo abelii* | 3 | F | 3/5/1979 | Zürich, CH | Hand/Parent |
| Hantu (HU) | *Pongo abelii* | 2 | M | 3/5/1981 | Zürich, CH | Parent |
| Radja (RJ) | *Pongo abelii* | 3 | F | 19/1/1973 | Zürich, CH | Parent |
| Timor (TR) | *Pongo abelii* | 2 | F | 23/11/1973 | Basel, CH | Parent |

#### Procedure

A baseline condition was implemented, in which a wooden hammer (25cm long, 8-10cm in diameter) was provided alongside shelled coula nuts, Brazil nuts and coconuts. No demonstrations on how to use the hammers to crack the nuts were provided before or during testing. The two *Pongo pygmaeus* received one coula nut per session for both individuals (one nut between both individuals was provided), five Brazil nuts each and one coconut each per session whilst the *Pongo abelii* received one coula nut and one coconut each and five Brazil nuts each per session. Each subject received one hammer. Initially no anvils were provided, as there were already objects inside the enclosure that could be used as anvils, such as logs with indentations that were in the orangutans’ outside enclosure. However, once it became clear that the *Pongo pygmaeus* preferred to manipulate the testing equipment in the indoor enclosure, an extra anvil location was created by chiselling a hole into the concrete of the *Pongo pygmaeus’* indoor enclosure flooring. The *Pongo abelii* were not provided with additional anvils (due to zoo management requirements, as their enclosure had just been renovated and they had more anvil-type objects that could be used). The nuts were scattered on the floor of the enclosure one hour before feeding. Following the testing protocols adopted at the time, once the testing session started, focal animals (defined as the ones that were currently holding a nut at the start of the session) were followed and observations reported at 10second intervals. In the cases when more that one individual was manipulating a nut, both subjects were followed. However, this happened rarely, and only a maximum of two individuals at a time manipulated a nut, making the combined focal follows manageable. Each session lasted one hour, and 15 sessions were carried out in total. Once the session ended, the keepers removed the hammers and the nuts from the enclosure. No video recording was made of the tests, as the video equipment was found to be too distracting to the subjects by the experimenter (MF). Observations were recorded by voice instead using a handheld tape recorder (unfortunately these tapes no longer exist).

## Results

The orangutans were observed practicing several different techniques to open the nuts (see Table 7 for an ethogram of these behaviours). Three individuals, one in each group (PG, AD & RJ), used the wooden hammer to crack open the coula nuts in the first testing session. In the subsequent second to seventh testing sessions, seven out of the total eight tested orangutans also demonstrated the same target hammering action (see Table 8 for the number of interactions with nuts with the hammering method observed across subjects). Therefore, at least three individuals spontaneously acquired nut-cracking with a hammer without having any demonstrators available. The only individual who did not demonstrate nut-cracking with a hammer was the youngest subject (HU), who rolled or threw away the nuts at the start of each session, and did not seem motivated to manipulate the nuts or the hammer. Although seven individuals attempted to crack nuts with the provided hammer, only three individuals (RS, RJ, TR) were successful in opening the nuts with this method, potentially due to the fact that not all individuals used an anvil to stabilise the nut. Most individuals only used the hammer to crack the coula nuts, as they were the only type of nut that required the hammer (both the Brazil nuts and the coconuts could be opened by biting or by hitting them on the floor or walls). However, two individuals (PG and RJ) also used the hammer on the coconut, of which only RJ was successful. PG was the only individual who was able to open the coula nuts with his teeth from the first session, and continued with this method for the duration of the study.

**Table 7:** Ethogram of behaviours directed towards the nuts observed in the Zürich study.

| Behaviour | Description | Successful for opening nut? |
| --- | --- | --- |
| Bite | The subject inserts the nut in its mouth and presses it between the teeth | Yes |
| Shake | The subject holds the nut in one or both hands and moves it back and forth quickly | No |
| Hit with hand/foot | The subject strikes the nut with either hands or feet, without using any tools | No |
| Press nut against teeth | After positioning the nut on the outer surface of the teeth, the subject presses it against the teeth either with the hands directly or by using a hard object such as the iron bars of the cage or wooden sticks | No |
| Scratch nut | The subject uses the nails to scratch the nut by damaging the surface of the shell | No |
| Rub nut | The subject moves the nut back and forth on a surface such as the ground or the walls | No |
| Beat against substrate | The subject holds the nut in one or both hands or feet and hits it against the floor or the walls or any other element in the enclosure | No |
| Press nut | The subject uses its weight or force on the nut without causing a sudden impact on the nut | No |
| Bounce nut | The subject throws or drops the nut onto the floor | No |
| Poke with branch | The subject pokes the shell of the nut with a branch | No |
| Wrapping nut | The subject wraps the nut with objects such as cardboard, wood wool or cotton bags | No |
| Hammering | The subject uses a hard object such as the wooden hammer to pound against the nut | Yes |

**Table 8:**
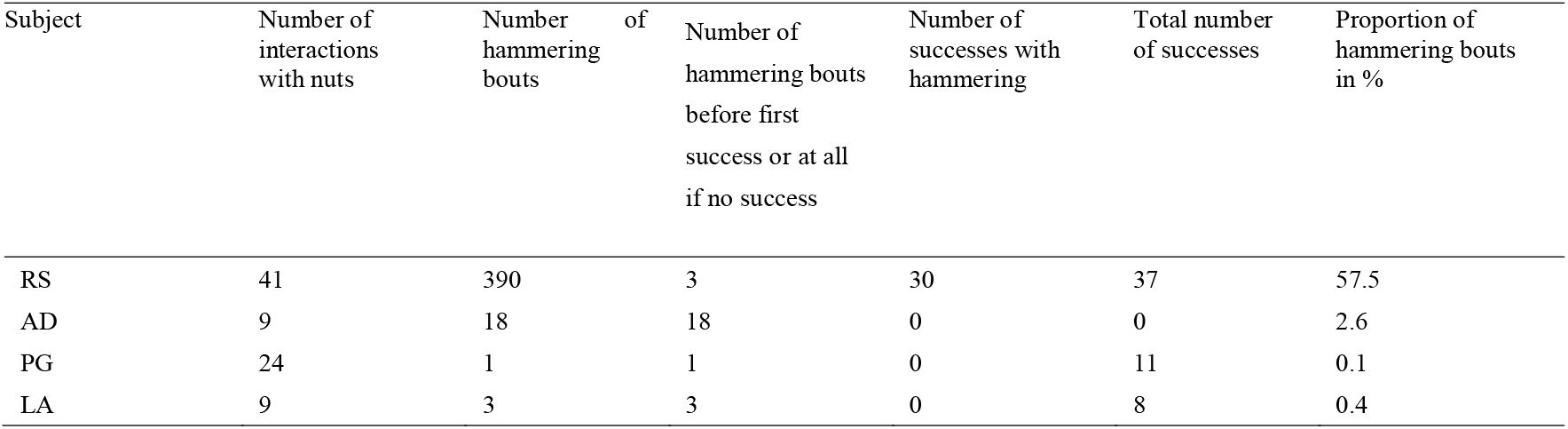
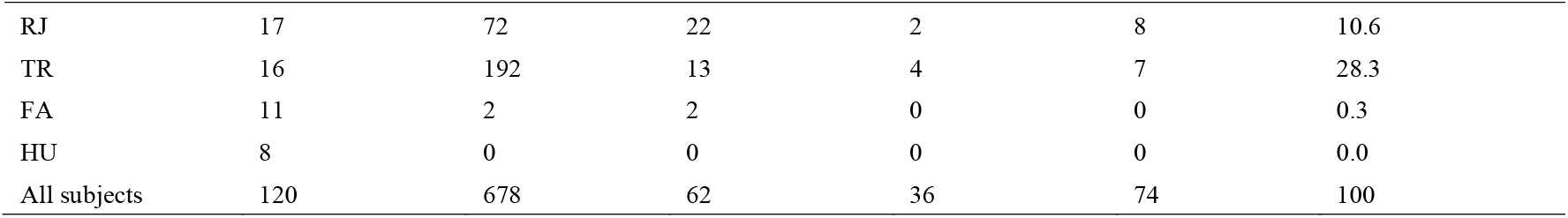
Total number of nut manipulation and hammering bouts in the Zürich zoo study.

## Discussion

Four naïve, unenculturated, captive orangutans spontaneously -without any copying or training opportunities-cracked nuts using a wooden hammer as a tool. This finding suggests that orangutans possess the individual ability to express nut-cracking in a similar way to wild chimpanzees, and that copying is not required for this behaviour to be expressed in this species. As orangutans do not crack nuts with tools in the wild, it is highly unlikely that any of the orangutans in Leipzig and Zürich studies had previously observed this behaviour from wild, or related, conspecifics. Furthermore, the subjects’ experiences in previous experiments, training and enrichment were discussed in detail at both institutions with their keepers, who assured us that the subjects had never been trained or demonstrated how to crack nuts, or even the hammering action, before testing. Although it is impossible to account for every minute of a subject’s life, keepers at both zoological institutions are heavily involved in the research studies and enrichment programs with their animals, making them the most reliable source on the animals’ knowledge of behavioural forms. Therefore, based on all the information available from the wild and the testing facilities, we are confident in assuming that all the orangutans tested here were naïve to nut cracking (and the fact that nut-cracking appeared across both institutions renders the naivety of the subjects even more likely).

This is not to say that the orangutans did not benefit from any previous experiences. Indeed, all of the subjects had prior experience with nuts, and therefore knew that force could be applied to the shells of the nuts to access the kernel inside. However, they only had experience with walnuts and hazelnuts, which can be opened relatively easily by using the teeth - even by juvenile orangutans -, and with coconuts, which can be opened by hitting them against the floor or walls. In contrast, the coula nuts provided in the Zürich study and the macadamia nuts provided in the Leipzig study require 2.8kn and 2.2kn to be opened, respectively (Visalberghi et al., 2008), which makes these alternative methods unviable. This, and the lack of suitable tool materials prior to our studies, may explain, at least in part, why none of the subjects in these studies had ever been observed using tools to crack nuts before.

### Candidate mechanisms behind nut-cracking in orangutans

The findings of both the Leipzig and the Zürich zoo studies suggest that nut-cracking with tools can be acquired individually by naïve orangutans without the need of copying, provided the availability of appropriate materials and the subjects’ motivation to do so, as well as, perhaps, a basic knowledge of the problem at hand (i.e., that shells often encase kernels and that force can be applied to break them open). We are not suggesting that nut-cracking is a hard-wired behaviour in orangutans. Although the ZLS hypothesis can also include such cases, ‘latent solutions’ is an umbrella term that subsumes behaviours spanning from highly genetically-determined behaviours to more learning-dependent behaviours, with the exception of copying-dependent behaviours (Tennie et al., in press). In the case of orangutan nut-cracking, there are several reasons to believe that more than instinct is at play. Firstly, despite the fact that several orangutans in our studies performed nut-cracking, long-term field studies with wild orangutans have not (yet) observed such behaviour (e.g., Krützen, Willems, & van Schaik, 2011, although note that this might also be due to a lack of need to crack open nuts). This suggests that nut-cracking is not (and is unlikely to have been) a critical behaviour for the species’ survival that may have been fixed in their genes throughout evolution (although note that some of the abilities required for nut-cracking, such as the manipulative tendency of the species, their hand affordance and some of their cognitive capacities may be more genetically determined). Secondly, there was variability in the performance of subjects in our studies. Despite experiencing the same conditions, not all the orangutans acquired the behaviour within the time frame given (although we acknowledge that motivation plays a role as well). Lastly, the orangutans in our studies demonstrated flexibility in their approach to the problem at hand – as an example, PD attempted several different methods to access the kernels before arriving at the target behavioural form of nut-cracking and, even after discovering the target behaviour, she did not then use it in every session. Perhaps most importantly, PD proved able to crack open nuts with a variety of tool use techniques.

If strong genetic predispositions and reliance on copying forms of social learning are excluded as explanations for the acquisition of this behaviour, a plausible alternative candidate mechanism is individual learning, supported by non-copying social learning. All apes demonstrate impressive abilities for such type of learning (see Tomasello & Call 1997; Whiten & Mesoudi, 2008 for an overview of these studies). Whilst individual learning abilities probably involve some genetic predispositions (see above), they also rely on cognitive skills that allow for considerable behavioural flexibility, including the capacity of finding different solutions to a given problem. One example of this flexibility is PD’s performance in this study. In the baseline, before the locked-anvil condition, PD used the provided large wooden block to crack open nuts, already demonstrating a similar tool use to wild chimpanzee nut-cracking but using a different tool and action. PD might have initially preferred to use the large block instead of the small wooden hammers as, although the former required more effort when being lifted due to its large weight (approx. 50kg vs. 2.4kg), it did not require the application of hitting force and speed to crack the nut, but could simply be part-lifted and/or rolled, and then dropped on top of the nuts. Moreover, the large block may have been easier to manipulate since its larger width required less precision when aiming to hit the nut than a hammer does. Once the large block was rendered inaccessible in the locked-anvil condition, however, PD flexibly switched her approach and used a hammer, demonstrating the behavioural form of nut-cracking similar to that observed in some wild chimpanzee populations (Biro et al., 2003; Boesch et al., 1994; Luncz & Boesch, 2014; Luncz et al., 2012). In brief, individual learning, alongside some genetic predispositions, non-copying social learning, and enhanced cognitive capacities that allow flexibility in the search for solutions to problems, may drive the acquisition of nut-cracking in orangutans.

### Potential explanations for the lack of reinnovation of the target behaviour by the remaining orangutans

None of the adult orangutans in the Leipzig zoo study used tools to crack macadamia nuts, and one juvenile individual in the Zürich zoo never attempted to use a hammer on coula nuts either. In the case of the Leipzig study, the adults were immediately and consistently successful in cracking open the nuts with their teeth, and continued doing so even after they were exposed to five sessions of live demonstrations of nut-cracking with a tool by PD. One explanation for the absence of nut-cracking with tools behaviour in these subjects could be precisely the fact that, as we observed, the adults were strong enough to bite through the shells of the nuts (note that, although macadamia nuts are hard, orangutans have a remarkable bite strength; Daegling, 2007), which might have rendered the use of a tool superfluous for them. Similarly, PG in the Zürich study was successful in opening the coula nuts with the bite method and he persisted with this strategy (he only used the hammer on coconuts). In contrast, the sub-adult PD attempted to bite nuts in the first session but failed, most likely because she had not yet developed the same jaw strength as the adults in the group. Therefore, PD may have been the only test subject motivated to find alternative methods to biting in order to access the kernels, including the use of tools to open the nuts. According to this explanation, if even harder nuts had been provided, rendering the bite methodology impossible, the adults in the group might have also acquired the tool-use behaviour. This hypothesis is supported by the findings of the Zürich study in which the harder coula nuts were provided and adults were reported to acquire the behaviour. An alternative to the jaw force explanation would be that age differences in inhibitory control (defined as the ability to stop a planned or on-going thought or action; Carlson & Wang, 2007; see also Albiach-Serrano, Guillén-Salazar, & Call, 2007; Amici, Aureli, & Call, 2008; Parrish et al., 2014) and functional fixedness (defined as “*the disinclination to use familiar objects or methods in novel ways*” Brosnan & Hopper, 2014, 2) encouraged PD to explore new solutions to the problem at hand while preventing the adults in the Leipzig study from finding the same solution. The fact that the only individual who did not attempt to use a hammer in the earlier Zürich study (HU) was the youngest member of the group does not support this explanation, although it may have been that this subject simply lacked the motivation to manipulate the nuts and the hammers or even to consume the kernels. Indeed, HU immediately rolled the nuts away when they were in his possession and did not attempt to retrieve them when other members of the group took them.

## Conclusion

Collectively, the results of our two studies demonstrate that individual learning (probably aided by several factors, such as genetic predispositions and cognitive capacities that allow to find solutions to problems in a flexible way) is sufficient for the acquisition of the behavioural form of nut-cracking in orangutans. Although this study did not find direct evidence for non-copying social learning increasing the frequency of the nut-cracking behaviour (as the older orangutans may have been fixed in their alternative, successful method of cracking open the nuts with their teeth in the Leipzig study), it is likely that, similar to other ape behaviours, non-copying variants of social learning can increase and stabilise the frequency of nut-cracking within populations – at least when these mechanisms apply across generations (see also the discussion in Moore, 2013). Therefore, the behavioural form of nut-cracking could, in principle, become another example of a SMR (Bandini & Tennie, 2017; 2019), for orangutans. Indeed, it is possible that orangutans may one day be found to express (or have expressed) nut-cracking behaviour in the wild - as a latent solution.

## Acknowledgements

The authors are very grateful to the Wolfgang Köhler Primate Research Centre (WKPRC), in Leipzig, Germany and to Zürich Zoo, in Switzerland, for providing the testing facilities. The authors also thank William Daniel Snyder, Li Li and David Boysen for coding and second coding, and Christian Nawroth, Yasmin Möbius and Daniela Hedwig for helpful discussions. We also thank Josep Call, Heinz Gretscher, Vincent Müller, Franziska Zemke, Josefine Kalbitz for support and feedback in the past, either directly or indirectly related to the current studies and Christophe Boesch for allowing access to the Zürich zoo thesis. MF is grateful to Hans Kummer for helping design the study, Christophe Boesch and Dr. G. Anzeberger for sourcing the coula nuts from the Ivory Coast and Bruno Schnyder for help during testing. EB and CT are supported by the Institutional Strategy of the University of Tübingen (Deutsche Forschungsgemeinschaft, ZUK 63). At the time of writing, CT was also supported by the ERC: This project has received funding from the European Research Council (ERC) under the European Union’s Horizon 2020 research and innovation programme (grant agreement n° 714658; STONECULT project).

